# The new role of carbonic anhydrases and bicarbonate in acclimatization and hypoxia stress responses in *Arabidopsis thaliana*

**DOI:** 10.1101/2024.08.13.607795

**Authors:** Alex Białas, Joanna Dąbrowska-Bronk, Piotr Gawroński, Stanisław Karpiński

## Abstract

Plant growth and stress responses largely depend on the chloroplast retrograde signaling. Stoichiometry of carbon dioxide assimilation and transpiration, efficiency of photosynthesis, and absorbed energy fate in photosystems between photochemistry, fluorescence and heat channels impact on the chloroplast retrograde signaling. Recent studies revealed that 22 kDa photosystem II protein (PsbS) and plant β carbonic anhydrases (βCAs), except their obvious functions, are also involved in regulation of plant stress responses. Obtained results suggest that simultaneous overexpression of *βCA1* and/or *βCA2* with *PsbS* genes leads to improved photoprotection, acclimation to variable light conditions, and water use efficiency. However, this was achieved on the costs of lower biomass gain in double and triple (oePsbSoeβCA1 and oePsbSoeβCA1βCA2, respectively) transgenic lines in comparison to Col-0, and *npq4-1* mutant. After bicarbonate fertilization we observed significant increase in biomass production in triple transgenic lines compared to oePsbS and *npq4-1* plants, but not to Col-0. Transcriptomic analysis revealed that bicarbonate treatment of double and triple transgenic lines specifically induced expression of genes and transcription factors related to hypoxia, freezing, drought, high light, and pathogen attack stress responses, contrary to other genotypes. Interestingly, expression of two of these transcription factors, DREB - CBF2 subfamily (A-1 of ERF/AP2), and BT2 were reduced in oe*PsbS* transgenic line. Our results suggest a novel regulatory role of βCAs and bicarbonate in the regulation of stress responses and plant productivity.

## Introduction

All aerobic life on Earth depends directly on oxygenic photosynthesis, which uses light energy to transform carbon dioxide (CO_2_) and water (H_2_O) into organic compounds and release oxygen (O_2_) to the atmosphere (Stirbet et al., 2019). Plants evolved the capacity to absorb more energy than they required to carry out electrical charge separation and photochemistry. This capacity in plants is required for induction of acclimatory and defense retrograde responses such as: variable light acclimation; Systemic Acquired Acclimation and Resistance (SAA and SAR) or Network Acquired Acclimation (NAA) (Mateo et al., 2004; Mühlenbock et al., 2008; Szechyńska-Hebda et al., 2010). Consequently, they have developed a mechanism to protect them from photodamage called non-photochemical quenching (NPQ, also known as qE), where the excess of harvested light energy is dissipated as heat and fluorescence (Müller et al., 2001). NPQ consists of a ΔpH and PsbS-dependent process (qE), photoinhibition (qI), state transition (qT), and zeaxanthin formation (qZ) (Baker 2008). Crucial for NPQ is a 22 kDa photosystem II PsbS protein belonging to the Chl a/b/xanthophyll-binding proteins [(light-harvesting complex (LHC)] superfamily which acts in the chloroplast as a lumen pH sensor (Fan et al., 2015). Underlying the photoprotection process is a mechanism for the response of the PsbS protein to the pH gradient (ΔpH) established between the stromal (pH ∼ 7.5) and lumenal (pH ∼ 5.5) sides of the thylakoid membrane under conditions of high or excess light (Takizawa et al., 2007; Górecka et al., 2020). PsbS can detect the acidification of the lumenal side and get activated. It was also shown that NPQ can be induced in the absence of PsbS, but at a much slower time or can be restored by enhanced ΔpH (Johnson and Ruban, 2010). Recently it was demonstrated that PsbS protein level influences the plastoquinone (PQ) pool redox status, thus electrical, ΔpH-, abscisic acid (ABA)-, and jasmonic acid (JA)-dependent retrograde signaling pathways from chloroplasts. These affect cellular light memory, acclimation, and defense responses (including cell death) induced by excess light episodes applied on low-light-acclimated plants, which resulted in cross-tolerance to pathogen attack and UV-C irradiation episodes.(Szechyńska-Hebda et al., 2010; Górecka et al., 2020). Recent studies showed that CAs activity is also important for NPQ and photosynthetic electron transport regulation (Dąbrowska et al., 2016; Shitov et al., 2018).

Nowadays many researchers wonder how to improve photosynthesis for C3 and C4 plants, thus biomass and seed yield during time of global warming. There are some suggested transgenic modifications to improve C3 plant’s photosynthesis such as: introducing the CO_2_ concentration mechanism from algae or C4 plants, optimizing the electron transport chain and photorespiration, improving recovery of NPQ and light absorption/conversion, and modifying Calvin Cycle enzymes (Price et al., 2013; Ort et al., 2015; Kromdijk et al., 2016). So far, to improve photoprotective mechanisms and biomass production, overexpression of genes coding for violaxanthin deepoxidase (VDE), PsbS and zeaxanthin epoxidase (ZEP) were provided into different plant species.

Overexpression of *VDE*, *PsbS*, and *ZEP* in tobacco plants showed improved NPQ and biomass production (Kromdijk et al., 2016). Conversely in *Arabidopsis* and potato plants, where overexpression of the same genes resulted in improved NPQ but reduced growth under fluctuating light (Garcia-Molina and Leister 2020; Lehretz et al., 2022). *Arabidopsis thaliana* overexpressed *PsbS* gene had a higher qE capacity during short-term high-light conditions. Transgenic plants were significantly more tolerant to transient photoinhibition expressed by having higher PSII maximal photochemical efficiency (*F*_v_/*F*_m_), contrary to the *npq4-1* mutant which lacks functional PsbS (Li et al. 2002, Górecka et al. 2020). These results indicate that increased photosynthetic productivity by, for example, improved relaxation of photoprotection depends not only on the species and its morphology, but is strictly related to the physiological and molecular basis of manipulated processes.

Plants have three families of CAs: α, β, and γ, where each family is represented by multiple isoforms in all species (Moroney et al., 2001). CAs ensure CO_2_ supply to phosphoenolpyruvate carboxylase (PEPC) in C4 and CAM plants, and RuBisCO in the Calvin-Benson cycle in C3 plants. It is also known that CAs in C3 plants are involved in stomatal movement, amino acid biosynthesis, lipid biosynthesis, and biotic/abiotic stress responses (Di Mario et al., 2017; Kolbe et al., 2018). *βCAs* are the most abundant in land plants, affect photosynthesis, and are found in specific subcellular compartments: *βCA1* is presented in the chloroplast, whereas *βCA2* is located in the cytosol, but the set and subcellular localization of CAs and its isoforms vary among species (Di Mario et al., 2017).

The latest studies shed new light on CAs involvement in biotic and abiotic stress responses such as drought, high light, high CO_2_, heat, high salinity, inorganic carbon deficit, excess bicarbonate, and response to pathogens (Wang et al., 2016; Kolbe et al., 2018; Hu et al., 2020; Polishchuk 2021). According to these results, CAs expression is regulated differently depending on the type and intensity of stress and on plant species. Our previous results indicated that CAs may detoxify HCO_3_^-^ and facilitate its use in photosynthesis (Dąbrowska et al., 2016). Lipid synthesis from free acetate is a minor pathway playing a role in the detoxification/reuse of ethanol, acetaldehyde, and acetate produced by fermentation under hypoxic conditions (Lin and Oliver 2008), and CAs may participate in these reactions along with acetyl coenzyme A synthetase. Recent research on CAs localization revealed that under stress conditions chloroplast βCAs and its isoforms could be translocated into the cytoplasm and nucleus, and binding the protein of the salicylic acid (SA) signaling cascade, but the physiological relevance of this phenomenon remains unclear.

Assuming the amelioration approaches of photosynthetic and photoprotective mechanisms in C3 plants dependences on several regulatory processes, we tested combined effect of *PsbS*, *βCA1* and *βCA2* overexpression in *Arabidopsis thaliana* on photosynthesis, photoprotection, high light and photoinhibition stress responses, and biomass production. Our results clearly indicate that overexpression of *PsbS*, *βCA1* and *βCA2* improved photoprotection and water use efficiency but decreased stomatal conductance and transpiration rate, thus the biomass yield has dropped. Increased yield and electron transport rate of PSI were also observed. Bicarbonate fertilization of oe*PsbS* plants precisely suppressed, while in the double and triple transgenic lines, oePsbSoeβCA1, oePsbSoeβCA1βCA2, precisely induced genes and transcription factors associated with response to hypoxia stress, freezing, drought, high light and pathogen attack. Our results strongly suggest a new retrograde signaling route regulated by βCAs, bicarbonate, and PsbS. The findings are discussed in the context of the potential of C3 plants to improve stress tolerance and plant productivity in the era of global warming.

## Results

### Simultaneous overexpression of *PsbS* and *βCAs* deregulates photosynthesis, photoprotection and biomass production

As described in Materials and Methods, 7 and 6 independent transgenic lines were obtained for double (oePsbSoeβCA1) and triple (oePsbSoeβCA1βCA2) constructs, respectively. Three independent transgenic lines with various overexpression of manipulated genes were chosen for further analysis. Representative expression level of these genes is presented (Supplementary Fig. S1A-D). As a reference genotypes, *Arabidopsis thaliana:* Col-0, *npq4-1* mutant, and oePsbS were chosen. To confirm the effect of *PsbS* and *βCAs* overexpression on photosynthetic productivity and photoprotection we measured the following parameters in 4-week-old plants: fresh and dry weight, NPQ, maximal quantum yield (QY_max_), total proton motive force (pmf) and pH gradient on thylakoid membrane (ΔpH), in plants after water and bicarbonate treatment (Fig. 1). Transgenic lines overexpressing *PsbS* and *βCAs* revealed significantly higher dry but not fresh biomass gain compared to *npq4-1* and oePsbS, but not to Col-0 control plants (Fig. 1A and B). Fresh and dry weight were reduced in *npq4-1* mutant after bicarbonate treatment in comparison to water. However, in oePsbS line only fresh weight was reduced in bicarbonate treatment in comparison to water. Greater dry mass productivity in double and triple transgenic lines after bicarbonate treatment in comparison to *npq1-4* and oePsbS lines correlates with decrease in NPQ value, but not with PSII QY_max_ changes (Fig. 1C and D). ECSt parameter (pmf) was similar in all analyzed lines after bicarbonate treatment, except *npq4-1* mutant line. Pmf was the highest in Col-0 and *npq4-1* under control conditions. (Fig. 1E). ΔpH was induced after bicarbonate fertilization in Col-0 and *npq4-1* plants, contrary to other genotypes (Fig. 1F). In general, bicarbonate fertilization improved total pmf (by its reduction) in all analyzed genotypes, but not in *npq4-1* plants (Fig. 1E). This correlates with higher fresh and dry biomass production in double and triple transgenic lines compared to oePsbS and *npq4-1* lines, but not to Col-0. Due to CO_2_ assimilation is driven by the light reaction products of photosynthesis – adenosine triphosphate (ATP) and nicotinamide adenine dinucleotide phosphate (NADP), the influence of β*CAs* overexpression on photosynthetic productivity was also carried in the greenhouse, under variable natural light conditions with bicarbonate fertilization (Supplementary Fig. S2). Significantly higher biomass production in triple (oePsbSoeβCA1βCA2) transgenic lines was observed, which may indicate an additive effect of *βCAs* overexpression or pivotal role of *βCA2* in HCO_3_^-^/CO_2_ transport, furthermore in fine-tuning regulation to the changing environmental conditions such as excess light and high temperature stresses.

**Figure 1.**
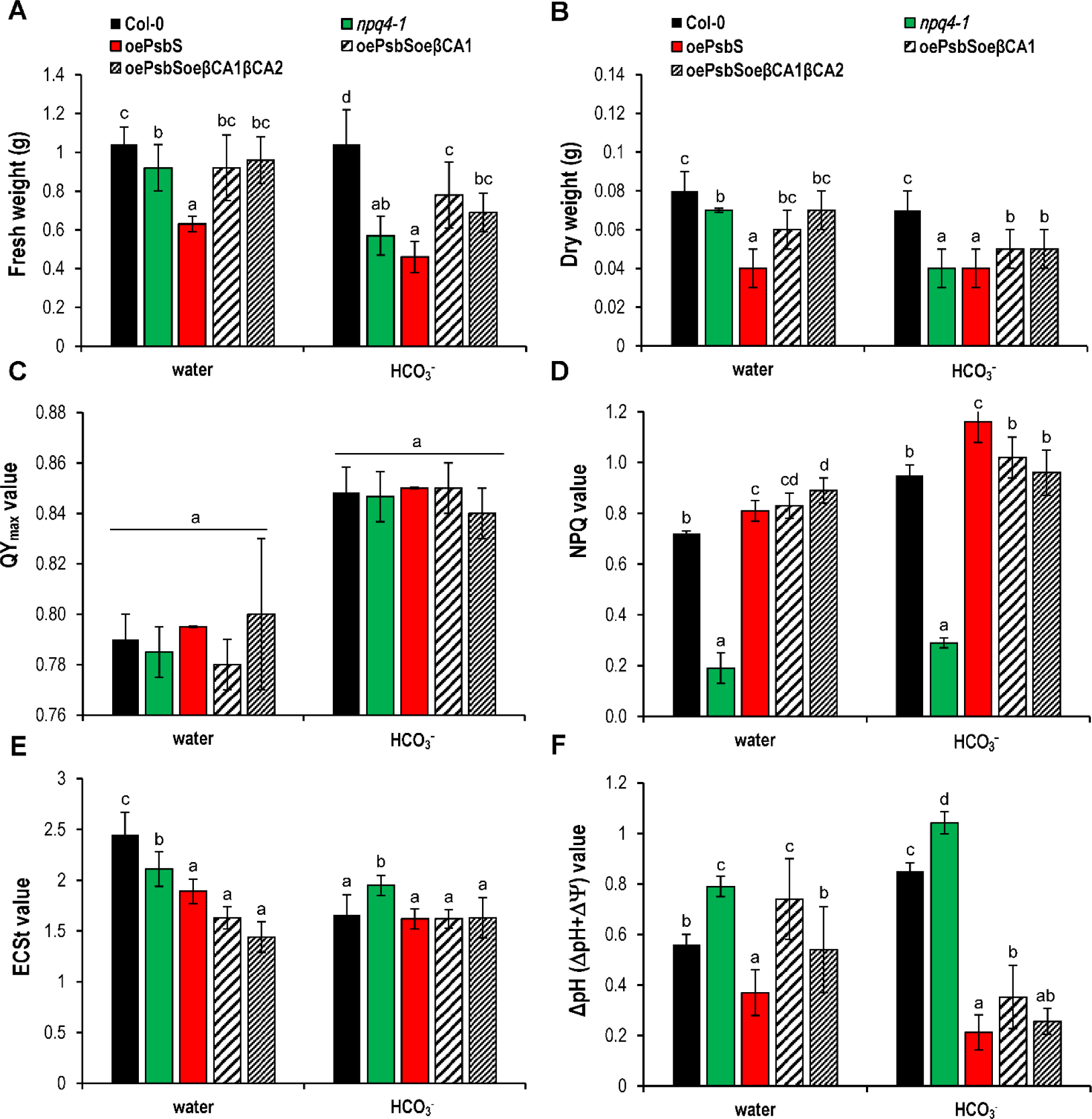
Overexpression of CA1 and CA2 influenced biomass production, NPQ and .ΔpH after water and bicarbonate supply. Effects on biomass productivity and photosynthetic parameters of bicarbonate fertilization of Col-0, *npq4-1,* oePsbS, double (oePsbSoe CA1) and triple (oePsbSoe CA1 CA2) overexpressing transgenic *Arabidopsis tha/iana* lines cultivated in ambient laboratory conditions. **(A)** Fresh and **(B)** dry weight of four-week-old plants. **(C)** Maximum yield of photosystem II (QY_maxl_ and **(D)** non-photochemical quenching (NPQ). **(E)** Total proton motive force and **(F)** npH parameters. Plants were cultivated in a growing chamber under normal light conditions 120 µE, fertilized with water or 3mM bicarbonate. One-way ANOVA Fisher’s least significant difference (LSD) procedure, 95% c.l, being used to estimate the difference between each pair of means. Data are shown as a mean ±SD (n=8 A-D, n=6 E,F). Results for double and triple transgenic lines are presented as an average value of three independent transgenic lines for each construct.

### Impact of β*CAs* on photoinhibition and Photosystem II photoprotection

QY_max_ and NPQ are sensitive indicators of changes in photosystem II that can lead to photoinhibition, degradation of D1 protein and photooxidative stress and in consequence cell death induction (Mateo et al., 2004). 3-(3,4-dichlorophenyl)-1,1-dimethylurea (DCMU) is a photosynthetic electron transport inhibitor that blocks reduction of the PQ pool by non-covalent binding to quinone B (Q_B_) side in PSII, and in consequence, slows down the oxidation of quinone A (Q_A_) and reduction of the PQ pool. DCMU treatment has been shown to strongly reduce NPQ, to induce cyclic electron transport (CET) and inhibits excess light-mediated induction of stomata closure and PQ pool redox status dependent retrograde signaling for light acclimatory, and immune defense responses (Mühlenbock et al., 2008; Szechyńska-Hebda et al., 2010). In our experiments, plants were treated with DCMU or exposed to high light (HL) to determine photoinhibition level in Col-0, transgenic and mutant plants. Our results revealed that PSII QY_max_ was significantly higher in double and triple transgenic lines after DCMU (Fig. 2A) and HL treatment (Fig. 2B). This indicates that overexpression of *βCAs* resulted in greater tolerance to moderate photoinhibition and HL stresses.

**Figure 2.**
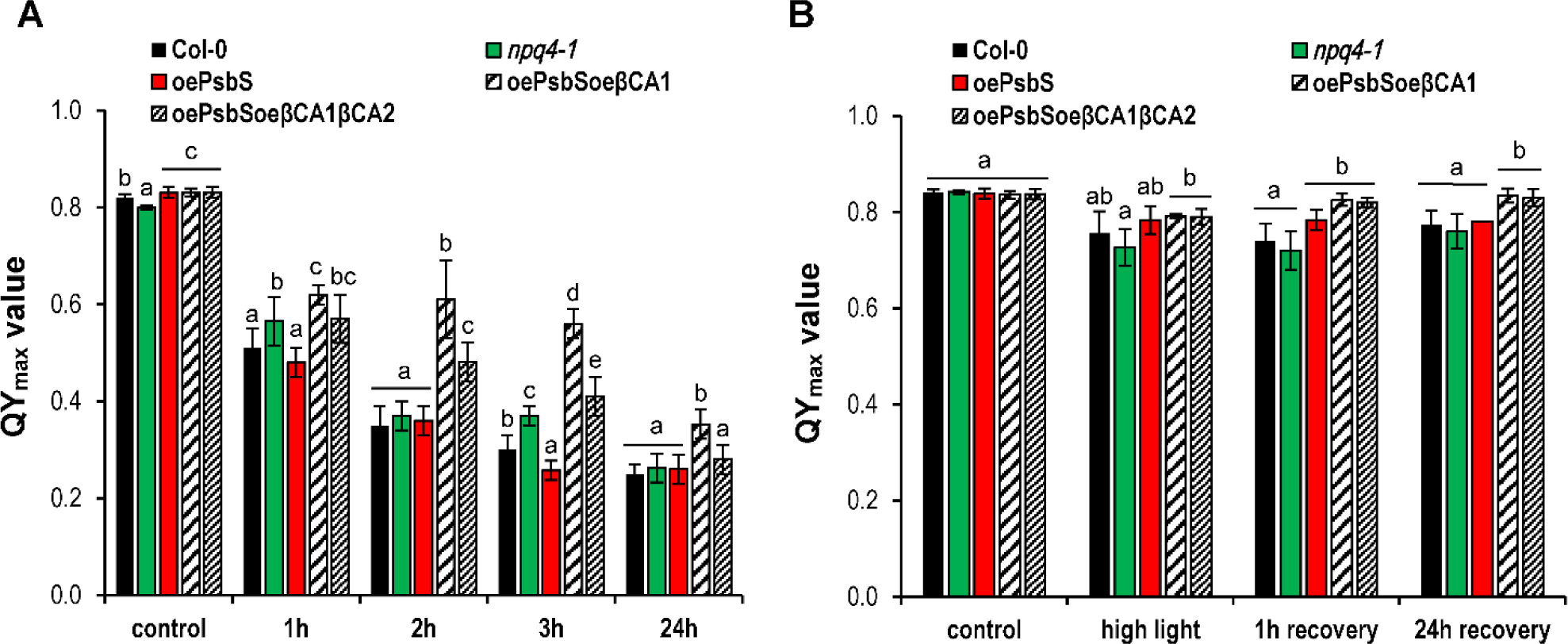
Overexpression of BCA1 and BCA2 led to improved photoprotection during and after photoinhibition. Effects on maximal photochemical efficiency of PSII (QY_max_) after 3(3,4-dichlorophenyl-)-1, 1-dimethylurea (DCUMU) and high light (HL) treatments of Col-0, *npq*4-1, oePsbS, double (oePsbSoeβCA1) and triple (oePsbSoeβCA1βCA2) overexpressing transgenic *Arabidopsis thaliana* lines cultivated in ambient laboratory conditions (**A**) QY*max* after 25 μM DCMU treatment and (**B**) HL (1000 μE) treatment. Four-week-old plants were cultivated in a growing chamber under laboratory ambient light conditions (120 μE) and exposed for 30 minutes episode of HL. One-way ANOVA Fisher’s least significant difference (LSD) procedure, 95% c.l, being used to estimate the differences between each pair of means. Data are shown as a mean ±SD (n=8). Results for double and triple transgenic lines are presented as an average value of three independent transgenic lines for each construct.

Knowing that DCMU and HL induce CET around PSI we simultaneously measured electron transport in PSI and PSII. Light response curves provide valuable information about the efficiency of photosystems in increasing irradiance and the use of photosynthetically active radiation (PAR) (Klughammer and Schreiber 1994). In light-dependent manner, active radiation absorbed by plants undergo three possible pathways: photochemical conversion yield (Y of PSI or PSII), dissipation of energy absorbed in excess in form of heat *via* the NPQ mechanism, and passively dissipated in form of heat/fluorescence, mainly due to closed PSII reaction centers Y(NO).

Immediately after the start of PAR illumination, the Y(II) values dropped from 0.8 to 0.6-0.5 for Col-0, *npq4-1* and oePsbS, and from 0.8 to 0.4-0.3 for double and triple transgenic lines, respectively (Fig. 3B), above 300 PAR there were no significant differences between genotypes. The lower Y(II) values in double and triple transgenic lines are primarily due to stronger regulated energy dissipation as heat, as reflected by the higher Y(NPQ) values (Fig. 3D). Y(NO) was similar in all genotypes at low PAR (Fig. 3F), but at higher PAR oePsbS had reduced, whereas *npq4-1* enhanced Y(NO), respectively. The variations in the ΔpH-dependent Y(NPQ) dominated the pattern of Y(II). The photochemical yield of PSI, Y(I), was higher in double and triple transgenic lines than in other genotypes (Fig. 3A). Due to increasing light intensity, >58 *µE*, higher limitation on the donor side of PSI, Y(ND), was observed in transgenic plants overexpressing βCAs (Fig. 3C), which was accompanied by lower Y(NA) (Fig. 3E). Overexpression of β*CAs* resulted in disrupted PSI:PSII stoichiometry, with a shift to PSI efficiency which is equivalent to running CET to protect donor and acceptor side.

**Figure 3.**
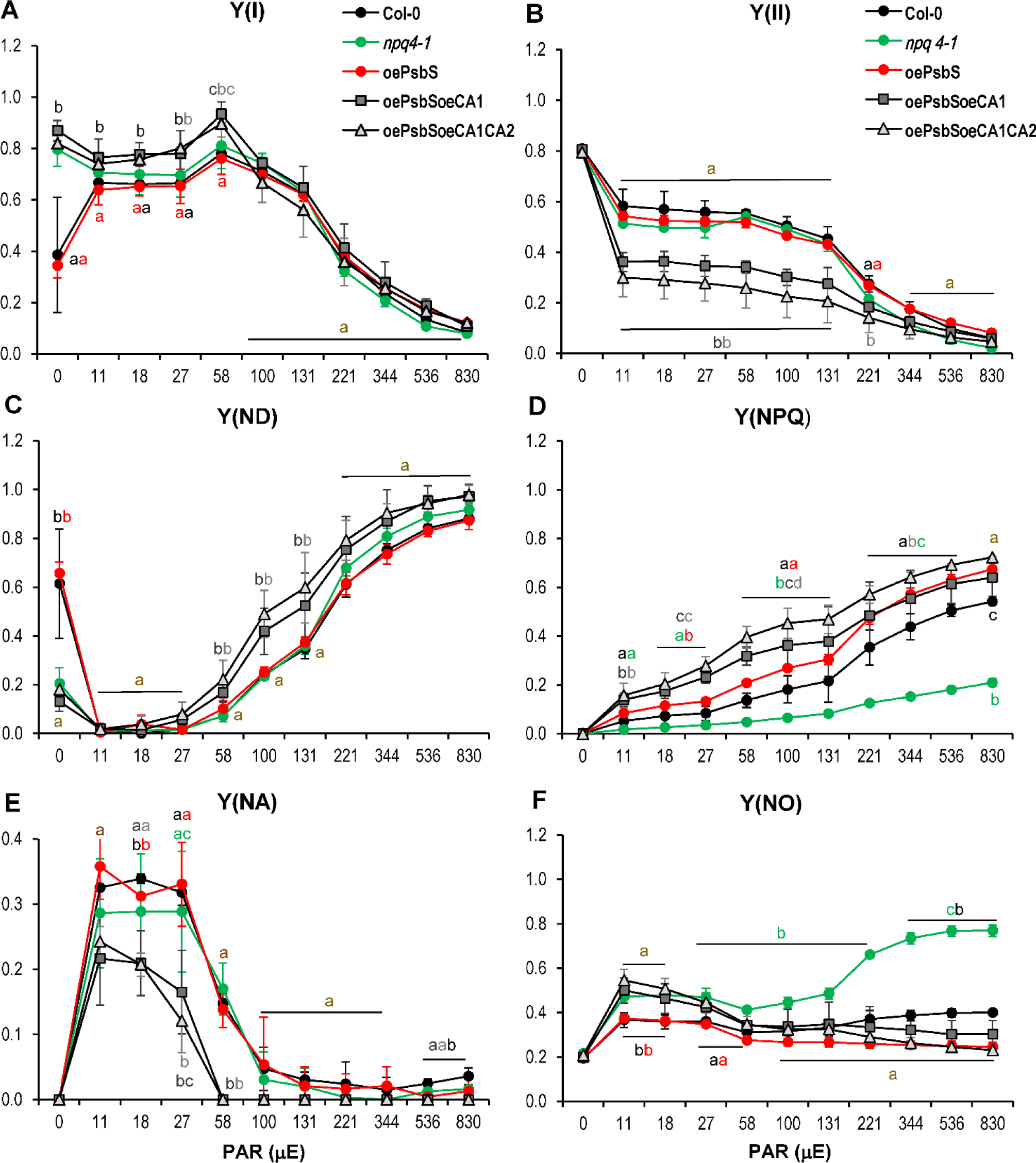
Overexpression of CA1 and CA2 influenced energy distribution and dissipation between photosystems. Light-response curves of the complementary quantum yields of Photosystem I and PSII of Col-0, *npq4-1,* oePsbS, double (oePsbSoe CA 1) and triple (oePsbSoe CA CA2) overexpressing transgenic *Arabidopsis thaliana* lines cultivated in ambient laboratory conditions **(A-F).** Four-week-old plants were cultivated in a growing chamber under normal light conditions of 120 µE Complementary quantum yields were measured at each PAR value (0-830). One-way ANOVA Fisher’s least significant difference (LSD) procedure, 95% c.l, being used to estimate the difference between each pair of means. Data are shown as a mean ±SD (n=8). Results for double and triple transgenic lines are presented as an average value of three independent transgenic lines for each construct. **Y{I)** - quantum yield of photochemical energy conversion in PSI; **Y(ND)** - quantum yield of non-photochemical energy dissipation due to donor side limitation in PSI; **Y(NA)** - quantum yield of non-photochemical energy dissipation due to acceptor side limitation in PSI; **Y(II)** - quantum yield of photochemical energy conversion in PSII; **Y(NPQ)** - non-photochemical quenching; **Y(NO)** - non-regulated energy dissipation.

### β*CAs* overexpression and bicarbonate treatment induce molecular response to photooxidative, drought, freezing and hypoxia stress

Double and triple transgenic lines significantly differed not only on physiological, but also molecular levels compared to Col-0, *npq4-1* and oePsbS. We aimed to check if there were any differences in genes expression patterns at the level of transcription. Transcriptomic analysis was performed using RNA sequencing (RNA-seq) on 4-week-old genotypes from control and bicarbonate-treated (3 mM) conditions. Fig. 4 presents all genes that are differentially expressed in bicarbonate fertilized genotypes compared to control water-treated plants. The analysis was conducted for each genotype separately (Fig. 4A). In Col-0, *npq4-1,* and oePsbS transcript levels were down-regulated after bicarbonate fertilization for most of genes. The situation was different in double and triple transgenic lines, where the vast majority of deregulated genes were induced (Fig. 4A). Out of 266 of all deregulated genes identified in double and triple transgenic lines, 73 genes were similarly deregulated in these lines (Fig. 4B). Gene ontology analysis for biological processes and molecular gene function (Fig. 4C and 4D, respectively) identified the two largest groups of genes involved in cellular response to hypoxia (over 20 genes induced out of 186 identified for hypoxia *in Arabidopsis thaliana* genome), and DNA binding transcription factors for drought, freezing, high light, high salinity, cold, and pathogen attack stress responses (Fig. 4E and 4F). Interestingly, the reduction in expression can be observed in 2 of these transcription factors after bicarbonate fertilization in oePsbS, indicating that overexpression of *βCAs* in oePsbS background reversed this deregulation (Fig. 4F). One of this gene encodes a member of the DREB subfamily A-1 of ERF/AP2 transcription factor family (CBF2) and second encodes protein (BT2) that is an essential component of the TAC1-mediated telomerase activation pathway. CBFs play important role in plant tolerance to abiotic stresses and activate the expression of other stress responsive genes in higher plants. In our analysis, among 266 genes differentially deregulated in double and triple transgenic lines after bicarbonate fertilization we identified 6 genes regulated by DREB TF *ERF/AP2* involved in freezing tolerance (*WRKY40*, *WRKY33*, *ZAT6*, *ZAT12*, *ERF105* and *CBF1*) (Fig. 4E and 4F, Supplemental Data Set 1).

**Figure 4.**
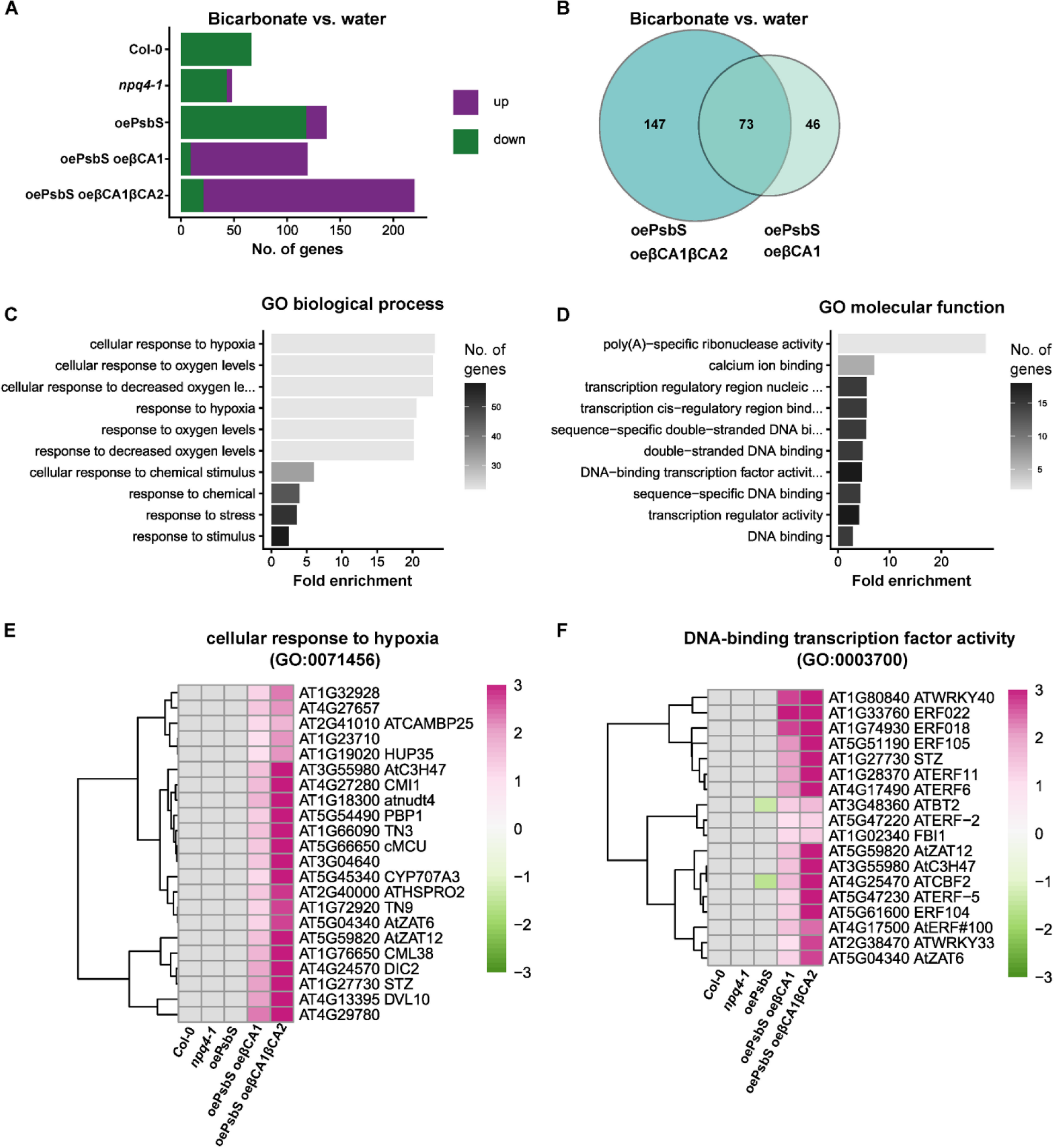
Differential gene expression pattern after bicarbonate *vs* water treatment of Col-0, *npq4-1,* oePsbS, double (oePsbSoe CA1) and triple (oePsbSoe CA CA2) overexpressing transgenic *Arabidopsis thaliana* lines cultivated in ambient laboratory conditions. **(A)** Diagram presents number of up- and down-regulated genes conversely regulated in analyzed genotypes after bicarbonate *vs* water treatment. **(B)** Venn diagram presents pools of genes regulated by bicarbonate in the oePsbSoe CA1 and oePsbSoe CA CA2 plants. Respectively, 147 and 46 genes were conversely regulated in these genotypes, 73 genes stayed in common. **(C)** and **(D)** Gene ontology analysis of oePsbSoe CA 1, oePsbSoe CA CA2 specific genes that are affected by bicarbonate treatment (in total, 73 genes were analyzed). **(E) and (F)** Clustering of genes specific to oePsbSoe CA 1, oePsbSoe CA CA2 plants belonging to “cellular response to hypoxia” and “DNA-binding transcription factor activity” Gene ontology groups.

## Discussion

In general, the fate of photons absorbed by photosystems can be divided into three different channels: for photochemistry, heat, and fluorescence. Efficiency increase in one process leads to decrease in the other two (Maxwell and Johnson, 2000). Significant mathematical correlation between high fluorescence, reduced stomatal conductance, and photosynthesis was also reported (Peak et al., 2004: Szechyńska-Hebda et al., 2010). Photosynthetic efficiency depends on many factors and processes in which the availability of HCO_3_^-^/CO_2_ and the dynamics of NPQ are pivotal, thus optimization of these processes seems to be crucial in crop productivity improvement. So far, scientists’ have been focused on modifying the NPQ process to improve plant productivity, but results were mostly contradictory, depending on the species and its physiology (Kromdijk et al., 2016; Hubbart et al., 2018; Garcia-Molina and Leister 2020; Lehretz et al., 2022).

During photosynthesis and its inhibition, reactive oxygen species (ROS) are produced mostly in chloroplasts and peroxisomes, which may cause photooxidative stress, but also play an important role in cellular redox retrograde signaling, regulation of plant growth, defense and light acclimation responses (Mateo et al., 2004; Mühlenbock et al., 2008; Szechyńska-Hebda et al., 2010; Czarnocka and Karpiński 2018). It is also widely recognized that PsbS serves as a crucial protein in the effective regulation of the fate of energy absorbed in excess. Previous studies demonstrated that the level of PsbS has an impact on the redox status of the PQ pool, thus on retrograde electrical and ROS signaling pathways (Górecka et al., 2020). It has been suggested that CAs participate in the regulation of chloroplast retrograde signaling and defense responses (Zhao et al., 2023). Based on available knowledge, we aimed to investigate whether βCAs interact with PsbS in regulation of the retrograde signaling from chloroplasts. Therefore we generated transgenic lines with overexpression of genes encoding βCA1, βCA2 and PsbS proteins.

In photosynthesis, light-driven electron transfer generates a pmf, consisting of an Δψ and a ΔpH (Kramer et al., 2003). The pmf drives ATP synthesis in thylakoids. While Mitchell’s chemiosmotic theory (1961) considers that Δψ and ΔpH are equivalent. Increased ΔpH triggers NPQ to protect photosystems from damage and regulates PQ oxidation. Conversely, increased Δψ can cause photodamage through singlet oxygen production (Davis et al., 2017). The PsbS protein and xanthophyll cycle regulate NPQ, but its induction and relaxation dynamics may sometimes negatively impact photosynthesis (Logan et al., 2008). It becomes obvious that fine-tuning of the NPQ process, their capacity, and dynamics, is principal in biomass improvement in plants. In our studies *βCA*s overexpression improved ΔpH (Fig. 1F) and significantly decreased the NPQ mechanism (Fig. 1D) according to the supply of bicarbonate ions, compared to oePsbS. Increased ΔpH and slightly enhanced NPQ positively influenced the supply of HCO_3_^-^/CO_2_, thus improved biomass production under ambient light conditions.

In our studies, overexpression of *βCA1* combined with overexpression of *PsbS* caused reduction in stomatal conductance (Supplementary Fig. S2B), thus in transpiration rate (Supplementary Fig. S2A), and improved WUE (Supplementary Fig. S2C) compared to oePsbS plants. The greatest effect was observed in triple (oePsbSoeβCA1βCA2) transgenic lines, where gas exchange parameters differ significantly from all control genotypes. However, CO_2_ assimilation stayed relatively unchanged in all tested lines (Supplementary Fig. S2D). Whereas CO_2_ assimilation was similar in the 500-800 ppm range. Photoinhibition (reduction of gas exchange) is reflected in higher NPQ but QY_max_ stayed in balance. This results explain reduced biomass in double and triple transgenic lines in ambient CO_2_ and water in comparison to Col-0, hence suggest that transgenic plants could be competitive in biomass production in elevated CO_2_ and Earth overheating.

CAs have diverse functions in plant cells and play an essential role in a wide range of biochemical and physiological processes - CO_2_ transport and sensing, respiration, pH regulation, amino acids synthesis, and acclimation to various environmental conditions. Earlier studies on *Arabidopsis* βCA1, 2, and 4 have demonstrated their involvement in abscisic acid (ABA)-independent CO_2_ signaling of stomatal closure in C3 and C4 plants, even under ambient CO_2_ level which affects gs and had a strong effect on increasing WUE under relatively unchanged carbon assimilation (Di Mario et al., 2017; Kolbe et al., 2018). Four distinct of βCAs - 1, 2, 3, and 4 were identified and their function was linked to lipid biosynthesis, antioxidant activity (CA1); providing bicarbonates to various metabolic pathways; amino acids synthesis (CA2, CA4), and providing a substrate for PEPC (CA3) (Dąbrowska-Bronk et al., 2016; Di Mario et al., 2016). In present studies overexpression of *βCAs* in oePsbS background improved biomass production after bicarbonate fertilization compared to oePsbS plants under ambient light, CO_2_, water, and temperature conditions (Fig. 1A).

Plants obtain carbon from both the atmosphere and water, where dissolved carbon concentrations are similar. HCO_3_^-^ enters plant roots, travels through xylem vessels to leaves, and after transformation by CAs to CO_2_ and water is assimilated with atmospheric CO_2_ (Poschenrieder et al., 2018). Moderate bicarbonate levels stimulate plant growth, but efficient conversion to carbohydrates in the Calvin-Benson-Basham cycle depends on light-driven ATP and NADPH production (Dąbrowska-Bronk et al., 2016). In oxygenic photosynthesizers, CO_2_ serves as the terminal electron acceptor for carbohydrate synthesis and regulates photosynthetic electron transport in PSII, which is responsible for light-induced charge separation and water oxidation. Research suggests that HCO_3_^-^ (CO_3_^-^), rather than CO_2_, may be crucial for PSII electron transport efficiency and the photo-assembly of its oxygen-evolving complex (Shevela et al., 2020; Shitov et al., 2018). This may explain slightly weaker biomass production for all tested *Arabidopsis* plants after bicarbonate fertilization, compared to plants irrigated with water under laboratory conditions. Experiments conducted in the greenhouse proved a higher impact of HCO_3_^-^ fertilization under variable light conditions on biomass production where triple overexpressing transgenic plants had significantly higher dry mass compared to oePsbS, *npq4-1* mutant, and oePsbSoeβCA1 plants (Supplementary Fig. S6A). This may indicate an additive nature of *βCA1* and *βCA2* or/and pivotal role of *βCA2* in binding inorganic carbon.

The role of CAs in photoprotection has been suggested recently (Dąbrowska-Bronk et al., 2016), where *Arabidopsis thaliana βcas* mutant plants revealed lower ion leakage and higher NPQ after bicarbonate supply. Moreover, in Col-0 plants, bicarbonate fertilization also improved the composition of photosynthetic pigments such as neoxanthin, violaxanthin, and carotenoids, which protect PSII and cells from, for example, high light stress. Additionally, the same role was proposed by Rudenko et al. (2018 and 2020), where α-*ca4* mutants had decreased major antenna proteins Lhcb1 and Lhcb2, but increased VAZ cycle components level. Overexpression of *βCA1* and *βCA1βCA2* in oePsbS background decreased D1, Lhcb1, and Lhcb2 transcript and protein levels (Supplementary Fig. S1 and Supplementary Fig. S3), but improved NPQ (Fig. 3D) and chlorophyll a/b ratio (Supplementary Fig. S4A) without influencing the carotenoids level (Supplementary Fig. S4B) under ambient light conditions. Therefore, oePsbSoeβCA1 and oePsbSoeβCA1βCA2 plants were more tolerant to high light stress in comparison to Col-0 and other analyzed genotypes (Fig. 2B).

DCMU is an effective photosynthetic inhibitor that binds to the Q_B_ binding site of PSII and inhibits photosynthetic electron transfer to the PQ pool, thus maintaining a reduced Q_A_ and oxidized PQ pool. Plants susceptible to DCMU have impaired photosynthetic efficiency, in consequence, lower biomass production (Gawroński et al., 2021). Our results indicate that double and triple transgenic lines had higher QY_max_ after DCMU treatment (Fig. 2A), thus the linear electron flow (LEF) between photosystems was blocked to a lesser extent than in other genotypes. On the other hand, the efficiency of electron transport in PSII (Supplementary Fig. S5), limitation of donor side Y(ND) (Fig. 3C) and photochemical quantum yield of PSII (Fig. 3B) were impaired, suggesting photoinhibition and reduced production of ATP and NADPH. As a result, alternative CET mechanisms were triggered which involve ferredoxin as an electron donor for PQ pool, the activation of NPQ, and regulation of the P_700_ redox state. When light levels are further enhanced and inhibit PSII with a substantial rise in NPQ, the flux through PSI continues, implying cyclic electron flow (CEF) involvement in photoprotection (Walker et al., 2014; Kono et al., 2014). It is known that CEF is inhibited in oePsbS plants, thus PSI is more vulnerable to damage during high-light stress. Experiments in the greenhouse (Supplementary Fig. S6) have proven our results that double and triple transgenic lines grew better under variable light conditions and had higher biomass compared to oePsbS plants.

Due to Earth overheating drought and floods are rising rapidly worldwide, which have a negative impact on agriculture. During waterlogging the plants are exposed to oxygen deprivation stresses like: hypoxia, anoxia, and reoxygenation. To cope with these stresses plants developed various strategies such as fermentation pathways, enzymatic and non-enzymatic antioxidant systems, cell death response like aerenchyma formation, and adventitious root growth to mitigate oxidative damage and enhance plant adaptation capacity during oxygen deprivation (Blokhina et al., 2003). Furthermore, the transition from low to high oxygen levels during re-oxygenation poses challenges due to excess ROS formation, prompting plants to activate mitochondrial retrograde signaling, chromatin modification, and alternative electron transport pathways to survive this stress (Jethva et al., 2022). Nevertheless, ROS are crucial in signaling pathways for acclimatory and defense responses in plants (Czarnocka and Karpiński, 2018), and activate response through cross-talk among various signal transduction pathways that encompass transcription factors. Our transcriptome analysis showed a very interesting and important regulatory phenomenon. Bicarbonate fertilization of double and triple transgenic lines identified 73 induced and co-regulated genes. Interestingly, for Col-0, *npq4-1*, and oePsbS after bicarbonate fertilization, transcription for all these genes, except two (DREB - CBF2 subfamily (*A-1 of ERF/AP2*) and *BT2*) in oePsbS, were not significantly altered in comparison to control (water treatment). Most of these deregulated genes are involved in hypoxia stress responses (Fig. 4C and 4E), and encode zinc-finger transcription factors (*ZAT6*, *ZAT12*, *ORA47*, *STZ*, *WRKY*, *TERF*, DREB - CBF2 subfamily (*A-1 of ERF/AP2*), *BT2* and *ERF*), that regulate the expression of downstream genes involved in various stress responses such as drought, cold, freezing, salinity, high light, heat, and pathogens (Davletova et al., 2005; Chen et al., 2016; Bolt et al., 2017). RNAseq also demonstrated a cross-talk between SA, ABA, ROS and ethylene (ET) in hypoxia stress response. Plants react to stress through complex processes involving ROS production, regulated by hormones like ET, JA, and SA (Gururani et al., 2015). ABA receptors and genes with ABRE and DRE elements increase under hypoxic conditions in plants (De Ollas et al., 2021; Correa Molinari et al., 2023). ET induces aerenchyma and ROS production, while SA is crucial for SAA, SAR, and cellular redox homeostasis. Overexpression of βCAs with PsbS, combined together with bicarbonate fertilization caused induction of *APX1*, *ZAT12* and *ZAT6* expression. This result strongly suggests that cellular βCAs activity and bicarbonate uptake are important in regulation of hypoxia stress responses. Interestingly, in tumors, hypoxia-induced changes are primarily regulated by HIF-1, which controls genes like CA9, crucial for pH modulation and CO_2_/proton movement across the cell membrane. During hypoxia in tumors, CA9 is overexpressed (Wykoff et al., 2001). In general, hypoxia reduces stomatal conductance and CO_2_ assimilation, thus photorespiration is induced, which strongly contributes to the induction of cell death. Double and triple overexpression lines have very high transcript levels of *βCA1* and/or *βCA2*, therefore respond better to hypoxia stress than the other genotypes. Likewise, high transcript levels of *βCA1* and/or *βCA2* combined with bicarbonate treatment induced DREB - CBF2 subfamily A-1 of *ERF/AP2*, which was inhibited in oePsbS plants, indicating that cellular levels of β*CAs* and oePsbS are crucial for fine-tuning of stress response. This implies that cellular CBF subfamily genes play a pivotal role in response to abiotic stresses in numerous plant species. Analysis of *Arabidopsis* mutants revealed that 134 genes are regulated by *CBF* genes in response to low-temperature stress (Jia et al., 2016). Our transcriptomic analysis revealed 6 genes regulated by DREB TF *ERF/AP2*, involved in freezing tolerance (*WRKY40*, *WRKY33*, *ZAT6*, *ZAT12*, *ERF105*, and *CBF1*). Our results suggests the existence of a new chloroplast retrograde regulatory hotspot and a new signaling pathway dependent on βCAs and PsbS protein relative levels and bicarbonate uptake (Fig. 5)

**Figure 5.**
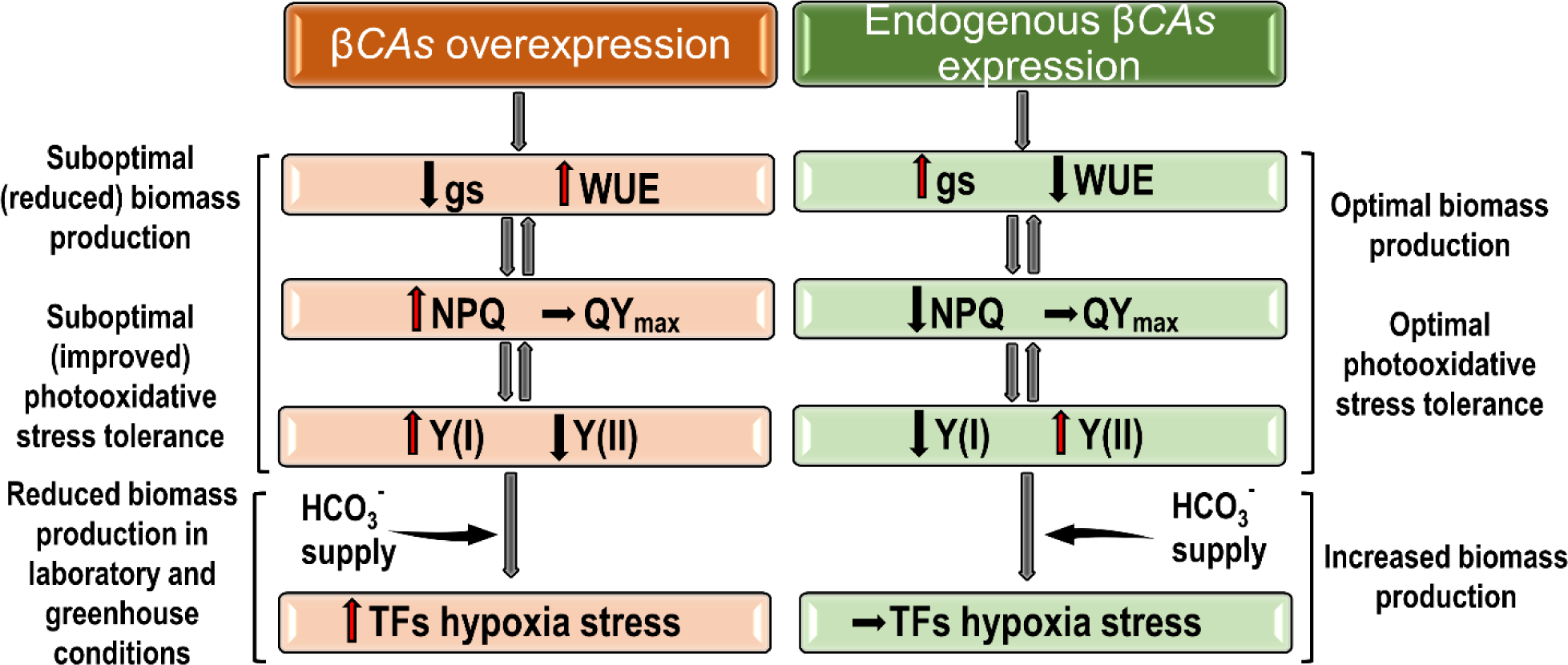
Scheme summarizing *Arabidopsis thaliana* response to *βCAs* overexpression. (described in the summary of the discussion)

To conclude, βCAs overexpression resulted in improved WUE and NPQ, as well as in the activation of transcription factors involved in various signaling pathways to defend against oxidative stress and cell death induced by hypoxia. In response to climate change, plants with *PsbS*, *βCA1*, and *βCA2* overexpression can be more adaptive to abiotic and biotic stresses such as hypoxia, photooxidation, cold, freezing, changing light conditions, high light, heat, salinity, and pathogens attack. It is worth noting that changes in the expression of CAs can give different effects depending on the species. Understanding these intricate molecular pathways is essential for improving plant tolerance and ensuring agricultural sustainability, thus further research is needed.

## Materials and methods

### 1. Plant material and growth condition General growth conditions

Wild type, mutants and transgenic plants were germinated on Jiffy pots (Jiffy Products). Pots were kept for 48 hr at 4°C and then placed in laboratory conditions – long-day photoperiod (16 hr light/8 hr dark) in 120 μE (NL), 80 μE (LL) and 800 μE (HL), and a temperature range of 20/22°C in a growth chamber (BDR16, Conviron). All experiments were conducted on *Arabidopsis thaliana* plants ecotype Columbia 0 (Col-0), *npq4-1-1* mutant - devoid of the *PsbS* gene (AT1G44575), plants overexpressing PsbS protein (oe*PsbS* Li et al. 2002), plants combine overexpressing *PsbS* and beta carbonic anhydrases 1 (*βCA1;* AT3G01500), and beta carbonic anhydrase 2 (*βCA2;* AT5G14740).

#### Bicarbonate fertilization

Wild type, mutant and transgenic plants were grown in normal light conditions (120 μE) and long photoperiod (16/8 h light/dark) in the growing chamber. Growth temperature was 21°C and humidity 50–60%. 2-week old plants, every two days were fertilized with 10 ml of 3 mM NaHCO_3_ (pH = 7). Control plants were treated with water at the same time and with the same volume.

#### Fresh and dry weight measurements

Fresh weight was collected from 4-week old plants (±1 mg). Then the rosettes were dried 24 hours at 150°C in a Hereus^®^ drier and weighed to establish dry weight (±0.1 mg).

### 2. Transgenic plants generation

*βCA1* (1044 bp) and *βCA2* (996 bp) coding sequences (The Arabidopsis Information Resource; www.arabidopsis.org) were amplified based on Arabidopsis Col-0 ecotype cDNA using Phusion II hot start high fidelity polymerase (Invitrogen) and cloned into pJET 1.2 vector according to manufacturer’s instruction (Thermo Fisher Scientific). Overexpression constructs for *βCA1* nad *βCA1βCA2* were generated using binary vectors pK7FWG2 and pK7m34GW2-8m21GW3D respectively, using „Gateway” technology (Karimi et al., 2002), according to manufacturer’s instruction (Invitrogen). Arabidopsis plants overexpressing PsbS protein were transformed with pK7FWG2::*βCA1* and pK7m34GW2-8m21GW3D::*βCA1βCA2* constructs based on *Agrobacterium*-mediated stable transformation using floral-dip method (Clough and Bent, 1998). Finally, 7 and 6 independent transgenic lines for oePsbS::oeβCA1 and oePsbS::oeβCA1βCA2 were obtained, respectively (T_3_ generation homozygous plants). Three of them, with the highest overexpression of *βCA1* and *βCA1βCA2* genes, were chosen for further analysis.

### 3. Gas exchange measurements

Gas exchange parameters such as net CO_2_ assimilation rate (A), transpiration (E), stomatal conductance (gs) and water use efficiency (WUE) were measured using CIRAS-3 Portable Photosynthesis System according to the manufacturer instruction (PP Systems, Amesbury, MA, USA). Eight leaves were collected per genotype. Measurements were performed in constant light intensity 120 μE. The temperature, CO_2_ concentration, and humidity in the measuring chamber was maintained at 25 °C, 400 ppm, and 60%, respectively.

### 4. DCMU and high light treatments

Leaf disks cut from 4-week old plants were treated with 25 μm of DCMU and with 1000 µE of white light for 30 minutes, then were kept in the growing chamber under ambient light between measurements. The maximum quantum yield of PSII (QY_max_) of leaf disks and rosettes was measured using FluorCam 800MF (Photon Systems Instruments, Czech Republic). As a control, we used leaf disks incubated in identical conditions without herbicide and plants non-treated with high light, respectively.

### 5. Chlorophyll *a* fluorescences

Chlorophyll a fluorescence was measured using an imaging chlorophyll fluorometer (FluorCam 800 MF, PSI, Czech Republic). Prior to measurements, the plants were dark adapted for 30 min to determine F_0_ and F_m_. Fluorescence parameters were determined according to the manufacturer’s instructions and previously described procedures (Baker, 2008). At least six plants were measured per genotype.

### 6. Analysis of quantum efficiency of PSI and PSII

Light response curves of the complementary quantum yields of PSI and PSII were measured at absorbance changes - 830 nm and 875 nm, with Dual-PAM-100 (Walz GmbH). At least four plants were measured per genotype. Analysis was performed for the fifth to seventh detached leaves prior dark adapted for 30 min. The parameters of PSI and PSII activity were calculated according to the manufacturer’s instructions and Niewiadomska et al., (2011).

### 7. Thylakoid delta pH measurements

Electrochromic shift (ECS) spectroscopic measurements were conducted using the DualPAM P515 module (Walz), which allows simultaneous measurements of the dual-beam 550–515 nm difference signal (Schreiber and Klughammer, 2008). Prior to measurements, plants were dark adapted for 30 min before the 200 μE actinic light (AL) was switched on. After 15 min of AL illumination, the ECS signal was measured for 2 min., and the dark relaxation of the ECS signal was measured for 50 sec. Immediately after the first measurement, the ECS signal was again recorded for 2 min after exposure to 660 μE AL, and for 50 sec. after switching it off. The protocol for ECS measurements of pmf components (ΔΨ and ΔpH) was based on a previously described method (Bailleul et al., 2010). Proton permeability was evaluated as gH^+^ = 1/τ after fitting the decay kinetics of ECS signal in the first 100 ms of dark relaxation with a single exponential. At least four plants were measured per genotype. Analysis was performed for the fifth to seventh detached leaves prior dark adapted for 30 min.

### 8. RNA isolation, sequencing, and real time RT-PCR analysis

Total RNA was extracted using the SpectrumTM Plant Total RNA kit (Sigma Aldrich) according to the manufacturer’s recommendations. Prior to qPCR analysis, RNA samples were subjected to DNase I treatment with DNAfree TM kit (Ambion, Applied Biosystems) for 30 min. Total RNA concentration was determined using UV–VIS spectrophotometer (Thermo Scientific, NanoDropTM). Reverse transcription reaction for cDNA synthesis was performed on 2μg of RNA using High Capacity cDNA Reverse Transcription Kit (Applied Biosystems) according to the manufacturer’s instructions. Real-time qPCR reaction was performed with 2.5 ng/ml of cDNA using 7500 Fast Real-Time PCR System (Applied Biosystems) according to the manufacturer’s instructions. Relative transcript levels were determined using 7500 Software (Applied Biosystems) by normalizing the threshold cycle number of each gene with *Arabidopsis thaliana* protein phosphatase 2A (*PP2A*) reference gene. Relative gene expression was calculated using the 2^-△△CT^ method (Rao et al., 2013). Each qPCR was performed for three biological samples and three technical repeats. Primer sequences used in this work are available in Table S1 (Supplementary data). According to transcriptome analysis RNA quality was tested on an ExpirionTM Automated Electrophoresis System (Bio-Rrad). Further, samples were processed at CeGaAT (Tübingen, Germany). Libraries were constructed using a TruSeq Stranded mRNA LT Sample Prep Kit (Illumina, San Diego, CA), and sequencing was conducted on the NovaSeq platform in (2 × 100 bp mode). The Arabidopsis genome sequence, annotation, and annotated sequence features were downloaded from TAIR Ensembl (Ensembl Plants, version 52 TAIR10 genome release). Reads were mapped to Arabidopsis thaliana cDNAs (Ensembl, TAIR 10, release 41) using Salmon software (Patro et al., 2017). Transcript-level abundances were imported into R and analyzed using the DESeq2 package (Love et al., 2014). Genes with log2 FC (fold change) value higher than 1 or lower than −1 and with adjusted p-value lower than .05 were considered as significantly affected. Gene ontology enrichment analysis of gene sets was conducted using the topGO package from Bioconductor.

### 9. Protein extraction and western blotting

Total protein was isolated with 4× Laemmli Buffer (Bio-Rad) from plant leaf tissue grinded in liquid N2. Double and triple transgenic lines are presented as a pooled material of three independent transgenic lines for each construct. Protein concentration in samples was quantified using RC DC Protein Assay (Bio-Rad). Samples were separated by Sodium Dodecyl Sulfate Polyacrylamide Gel Electrophoresis (SDS-PAGE) on 10% acrylamide gel, each well was loaded with 20 μg of total protein. After separation samples were transferred from gel to Immobilon P PVDF membrane (Merck, Germany) by semi-dry transfer, then blocked with 1% NFDM (Bio-Rad) o/n at 4℃. The dilutions of primary and secondary antibodies are presented in Table S2 (Supplementary data). Chemiluminescence signal was detected using Pierce ECL Plus Western Blotting Substrate (Thermo Fisher Scientific) and ChemiDoc Imaging Systems (Bio-Rad). To confirm that wells were loaded with the same amount of protein, analogical gels were stained with QC Colloidal Coomassie Stain (Bio-Rad).

### 10. Antioxidant content

4-week old plants were grinded in liquid N2, then approximately 10-15 mg of tissue was suspended in pure methanol. Absorbance was measured using Multiskan GO (Thermo Fisher Scientific) as described before by Gawroński et al. (2021). Calculations were performed as reported previously by Sumanta et al. (2014).

